# Major Histocompatibility Complex class I heavy chains localize in both cytoplasmic and nuclear compartment

**DOI:** 10.1101/2022.09.13.507738

**Authors:** Maria Gómez-Herranz, Alicja Dziadosz, Sara Mikac, Michał Rychłowski, Robin Fahraeus, Natalia Marek-Trzonkowska, Elżbieta Chruściel, Zuzanna Urban-Wójciuk, Ines Papak, Łukasz Arcimowicz, Tomasz Marjanski, Witold Rzyman, Alicja Sznarkowska

## Abstract

The Major Histocompatibility Complex class I (MHC-I) molecules present antigenic peptides (AP) to CD8^+^ T cells for self versus non-self recognition. Loading of AP on MHC-I takes place in the endoplasmic reticulum (ER), upon shuttling of cytoplasmic AP substrates to the ER. Understanding of this process has been influenced by the view that MHC-I antigens are produced from the proteasomal degradation of cellular proteins. Recent observations on the intronic and untranslated region-derived peptides as well as on the non-AUG translation products presented on the MHC-I open the possibility that antigenic peptides can derive from pre-spliced mRNAs translated in the nuclear compartment. In this brief report, we show that a fraction of human MHC-I molecules (human leukocyte antigens type A, HLA-A) is present in the nuclei of cells, in the proximity of histone H2B. With this finding, we hope to initiate a new direction of research on the nuclear role of MHC-I and ask whether the loading of antigens can take place in the nuclear compartment.

## Introduction

Major Histocompatibility Class I (MHC-I) molecules present antigenic peptides of 8-10 amino acids on the surface of every nucleated cell for recognition by the CD8^+^ T cells [1,2]. Recognition of the non-self antigen, coming eg. from viruses or genomic alterations acquired during cellular transformation, activates T cells and leads to the destruction of infected or tumor cells [3]. It also accounts for the graft-versus-host disease after organ transplantation [4]. Therefore, the optimal function of the MHC-I system is fundamental to the fitness and survival of the organism.

In humans, the MHC-I genes, referred to as human leukocyte antigen (HLA) class I genes, are ubiquitously expressed and highly polymorphic [5]. They are subdivided into classical (HLA-A, HLA-B and HLA-C) and non-classical (HLA-E, HLA-F and HLA-G) genes and encode heavy chains of MHC-I molecules [6]. Upon assembly with a light chain (beta-2-microglobulin) in the ER lumen, MHC-I is glycosylated and rapidly recruited to the peptide loading complex (PLC) which facilitates loading of antigenic peptide on the MHC-I [7,8]. PLC consists of the MHC-I specific chaperon - tapasin, transporter associated with antigen processing (TAP), thiol oxidoreductase ERp57 and lectin-like chaperone calreticulin. The current understanding of the loading of antigens on MHC-I has been influenced by the view on their origin from the products of proteasomal degradation of full-length proteins in the cytoplasm. Consequently, AP substrates need to be transported through the ER membrane by TAP and trimmed by the resident aminopeptidases to peptides of 8-10 amino acids length [9]. After an antigenic peptide is bound to MHC-I (pMHC), the pMHC complex dissociates from PLC, proceeds through Golgi where it is sorted to the cell surface and gets anchored on the cellular membrane [10,11]. Upon ubiquitin modification and changes in their conformation, pMHC are recycled back, guided to multivesicular bodies (MVB) and directed to the lysosomes for degradation [12].

In this report, we demonstrate that MHC-I heavy chains localize in the nucleus of normal and cancer cells in the proximity of histone H2B.

## Results

To evaluate cellular localization of HLA-A we have used a panel of lung cells comprised of the non-small cell lung cancer (NSCLC) cell lines A549 and RERF, two primary cell lines derived from NSCLC patients (NSCLC-1 and NSCLC-2) and normal lung fibroblasts (NLF).

Immunofluorescence staining with anti-HLA-A antibody showed that HLA-A heavy chains dominantly localized in the nuclei of all tested cells (Fig. 1A). HLA-A was also observed in the cytoplasm, particularly cytoplasmic granules most probably corresponding to the MHC-I transported via ER and Golgi apparatus. Control staining with secondary antibodies proved that nuclear signal comes from anti-HLA-A antibodies (Fig. 1A, right panel). Cellular fractionation followed by HLA-A detection in Western blot confirmed that HLA-A localized to both cytosolic and nuclear fractions in all cell lines (Fig. 1B) and HLA-A knockdown with a pool of small-interfering RNAs validated the specificity of anti-HLA-A antibodies used in the study (Fig. 1C). Immune fluorescence with another anti-HLA-A antibody validated nuclear localization of HLA-As in A549 and RERF cells (Fig. 1D). Ectopically expressed HLA-A also localized in the nuclear compartment, as observed with immunofluorescence (Fig. 1E) and cellular fractionation (Fig. 1F). Moreover, endogenous HLA-A localized in the nucleus in proximity to histone H2B as detected with the *in situ* proximity ligation assay (PLA) (Fig. 1G and H).

**Figure 1.**
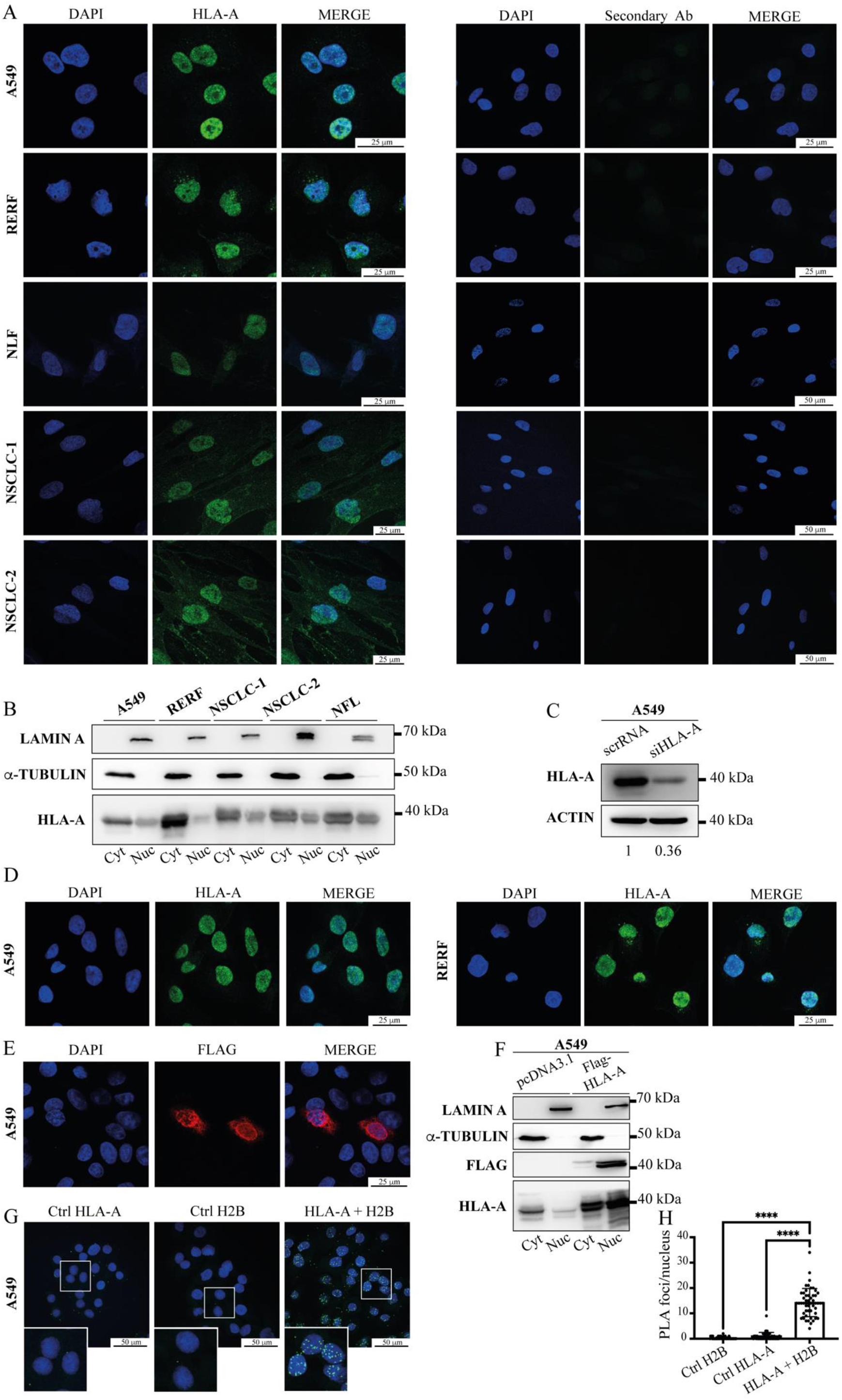
Evaluation of the nuclear HLA-A fraction in lung cells. **A)** Immunofluorescence-based detection of HLA-A in lung cells. Cells were fixed with 4% paraformaldehyde (PFA) and permeabilized with 0.2% Triton X-100. Cells were stained with anti-HLA-A antibody (Abcam, Cat #EP1395Y) and DAPI for nucleus localization (left panel) or just with secondaries antibodies (Secondary Ab) and DAPI in the right panel. Representative images are shown for five cell lines: A549, RERF, NLF, NSCLC-1 and NSCLC-2. Combination of both channels is represented in the merge column. Scale bar is 25 or 50 μm. **B)** Cellular fractionation of endogenous HLA-A in A549, RERF, NSCLC-1, NSCLC-2 and NFL cells. Cytoplasmic and nuclear fractions correspond to Cyt and Nuc lanes, respectively. **C)** Knockdown of *HLA-A* with specific small-interfering RNAs in A549 cells. Relative band intensities are presented below the blot. **D)** Immunofluorescence-based detection of HLA-A in A549 and RERF cells. Cells were fixed with 4% paraformaldehyde (PFA), permeabilized with 0.2% Triton X-100 and stained with anti-HLA-A antibody (Invitrogen, Cat #PA5-29911) and DAPI for nucleus localization. Scale bar is 25 μm. **E)** Immunofluorescence with anti-Flag antibody in A549 cancer cells transfected with Flag-HLA-A for 48 h. Nuclei are stained with DAPI (in blue). Combination of both channels is represented in merge. Scale bar is 25 μm. **F)** Cellular fractionation in A549 cancer cells transfected with Flag-HLA-A or pcDNA3.1 (negative control) plasmids for 48 h. HLA-A was detected with anti-Flag and anti-HLA-A antibodies. Cytoplasmic and nuclear fractions correspond to Cyt and Nuc lanes, respectively. **G)** Proximity ligation assay (PLA) in A549 cancer cells showing the detection of endogenous protein-protein association between HLA-A and the nuclear protein histone H2B. Nuclei are stained with DAPI (in blue). Negative controls employ anti-HLA-A antibody (Ctrl HLA-A) or anti-Histone H2B antibody (Ctrl H2B) only. The total number of nuclear foci per cell (in green) increases drastically with the detection of HLA-A and histone H2B together (HLA-A + H2B). Magnifications of selected areas are shown at the bottom left of each image. Scale bar is 50 μm. **H)** Representation of the relative quantification (50 cells per condition) of nuclear green fluorescent foci per cell with mean ± standard deviation. Statistical analysis was performed with pairwise two-tailed T-test (**** p< 0.0001).

## Discussion

This study shows that HLA-A localizes to the nucleus of cancer and normal lung cells and is found in the proximity of histone H2B. The presence of MHC-I in the nucleus cannot be explained based on the current understanding of the MHC-I pathway, according to which the synthesis and loading of MHC-I takes place in the ER lumen. An origin of antigenic peptides from a degradation of full-length proteins has been challenged by a number of observations, including the rapid presentation of viral antigens in a timeframe shorter than required for a turnover of a corresponding protein [13], identification of peptides coming from “non-coding” mRNA sequences such as introns [14], 3’ and 5’ untranslated regions [15] or long non-coding RNAs [16] and presentation of antigens generated by the non-AUG translation [17,18]. Finally, fusing the antigenic peptide to the IκBα sequence and inducing IκBα degradation did not show to have any impact on the presentation of the fused antigen, despite a giant increase in degradation substrates [14]. Recent studies show that DNA damage induces presentation of peptides coming from the pioneer round of translation in the nucleus [19]. Altogether undermined degradation of mature proteins as a source of AP. Simultaneously, antigenic peptides encoded within intron sequences were translated in the nuclear compartment from pre-spliced mRNAs, presented on the cell surface and activated corresponding CD8^+^ T cells [14,20]. These observations point to the conclusion that synthesis of antigenic peptides is separated in time and space from the synthesis of full-length proteins.

The fact that the presence of MHC-I molecules in the nucleus has not been spotted for that long might have its reasons. First of all, ‘what happens where’ during the class I presentation, was assumed to be known, thus *in situ* localization of MHC-I has just not been studied. A recent focus was more on finding out what are the peptides presented on MHC-I molecules for T cells immunosurveillance [21]. These studies rely on the w6/32 antibody that recognises a conformational epitope of an assembled MHC-I molecule, but w6/32 cannot really be used in fixed (cross-linked) cells, in which the conformational epitope might have been perturbed.

The presence of HLA-A in the proximity of histone seems to be associated with the nuclear function of HLA-A. Histones allow for spatial organization of chromatin and the accessibility of the DNA to be transcribed. Their association with HLA-A indicates that HLA-A might participate in the remodeling of the chromatin. The fact that the ectopically expressed HLA-A translocates to the nucleus provides a tool to further study its nuclear function with techniques such as immunoprecipitation and mass-spectrometry based analysis of binding partners. We hope that this observation will encourage research in this direction, as it seems that there is still much to understand about the class I pathway.

## Materials and Methods

*The detailed description of Materials and Methods used in this study is provided in the supplementary information*.

## Acknowledgments

The study was supported by “International Centre for Cancer Vaccine Science” that is carried out within the International Research Agendas Programme of the Foundation for Polish Science co-financed by the European Union under the European Regional Development Fund (MAB/2017/3), Polish National Science Centre PRELUDIUM9 grant No. 2015/17/N/NZ3/03773 and UGrants Advanced No. 533-0B00-GA08-22.

## Supplementary Information for

### This PDF file includes

Material and Methods

## Materials and Methods

### Cell lines

Non-small cell lung cancer cell lines A549, RERF-LC-AI and normal lung fibroblasts (NLF) were purchased from RIKEN BRC Cell Bank (Tsukuba, Ibaraki, Japan). A549 and RERF-LC-AI (referred in the text as RERF) were cultured in Dulbecco’s modified Eagle’s medium (Gibco, Thermo Fisher Scientific), with 8% of Fetal Bovine Serum (Gibco, Thermo Fisher Scientific) and 1% of Penicillin-Streptomycin (10 000 U/mL, Gibco, Thermo Fisher Scientific). Patient derived NSCLC-1 and NSCLC-2 cells and NLF cells were cultured in Ham’s F-12 Nutrient Mix medium (Gibco, Thermo Fisher Scientific) with 8% of Fetal Bovine Serum (Gibco, Thermo Fisher Scientific) and 1% of Penicillin-Streptomycin (10 000 U/mL, Gibco, Thermo Fisher Scientific). Cells were maintained at 37°C under humidified conditions with 5% CO_2_.

### Isolation of NSCLC cells and depletion of CD45+ cells

Cancer samples (0.25-1 g) were obtained from patients diagnosed with non-small cell lung cancer (NSCLC). Patients were admitted to the Clinic of Thoracic Surgery of University Clinical Centre of Medical University of Gdansk (MUG) for surgical resection of the primary tumor. They had not been previously treated with any anti-cancer therapy and no metastases were detected. Samples were cut into 1-2 mm fragments and washed 3 times with PBS to remove contaminating debris and erythrocytes. Subsequently, digestion solution (DS) in 1:1 of DS (ml): tissue mass (g) ratio was added. Tissue was gently agitated at 37°C until single cell suspension was obtained. DS contained 5% collagenase type I (Sigma-Aldrich; 5mg/ml), 19% of PBS and 1% of Fetal Bovine Serum (FBS). The collagenase was inactivated with an equal volume of 10% FBS supplemented LG-DMEM medium (PAA), followed by filtration of the resulting cell suspension through a 100 µm nylon cell strainer (Falcon). Subsequently, the cells were washed with PBS. Then, an erythrocyte lysis buffer was used to remove erythrocytes (10 min incubation at room temperature). After subsequent centrifugation (600 g, 10 min) the cells were washed with 4% FBS supplemented PBS and subjected for tumor infiltrating leukocyte depletion. For this purpose, positive immunomagnetic selection of CD45^+^ cells was performed with EasySep Human CD45 Depletion Kit II (StemCell Technologies, Canada) according to the manufacturer instructions (post-isolation purity 96-99%). Thus, untouched NSCLC and lung-derived cells were isolated. Cells were plated into 75 cm_2_ culture flasks in TumorPlus263 medium without serum. Establishment of the primary cell line was confirmed with flow cytometry (FAP-CK19^+^ phenotype; FACS Aria Fusion, BD Biosciences, USA) and when continuous proliferation of the cultured cells was observed after 2 month expansion in vitro. Cells that fulfilled both criteria were used for the experiments.

### Lipid-mediated transfection

A549 cells were seeded in the 6-well plates, 200 000 cells/well. 24 h after seeding, cells were transfected with siRNA-A (ON-TARGET plusTM Control Pool, DharmaconTM, referred in text as scrRNA, 25 nM), and HLA-A RNAs (siRNA, ON-TARGET plus TM SMART pool, DharmaconTM) in concentration of 25 nM, with 4 μl/well of Lipofectamine 3000 reagent (Thermo Fisher Scientific), according to manufacturer’s instructions. Following Lipofectamine 3000 protocol, A549 cells were transfected with Flag-tagged HLA-A or pcDNA3.1 plasmids to study the ectopic expression of HLA-A. Western blot and immunofluorescence were performed 48 h after transfection.

### Cellular fractionation

Searation of nuclei from cytoplasm was performed according to the REAP method described by Suzuki et al. [1]. Briefly, 2 000 000 cells were resuspended in 200 µL of ice-cold 0.1 % NP-40 in PBS by gentle pipetting and centrifuged at 500 g for 10 sec. Supernatant (cytosolic fraction) was collected and the Laemmli buffer was added to a final concentration of 1x. Pellets containing nuclei were washed with 200 µL of 0.1 % NP-40 in PBS, centrifuged at 500 g for 10 sec, resuspended in 200 µL 1x Laemmli buffer and sonicated 15 min. Samples were boiled for 10 min and analyzed by Western blot.

### Western blot

Total protein was acquired by lysing cells in the RIPA buffer. Proteins were electrophoretically separated via 8% SDS-PAGE and transferred to a nitrocellulose blotting membrane (Amersham Protran®). To block the membranes, 5% non-fat milk in Tris-buffered saline was applied at room temperature for half an hour. Membranes were subsequently incubated overnight with primary antibodies: rabbit mAb anti-HLA-A (cat. no. ab52922; Abcam); mouse mAb anti-Flag (cat. no. MA1-91878; Invitrogen); mouse mAb anti-α-Tubulin (cat. no. 3873; Cell Signaling Technology); mouse mAb anti-Lamin A (cat. no. 518013; Santa Cruz Biotechnology) or mouse mAb anti-β-Actin (cat. no. A2228; Sigma-Aldrich) in blocking buffer at 1:500 or 1:1000 dilutions at 4°C. Subsequently, membranes were washed three times in TBST followed by incubation for 1 h with HRP-labeled goat anti-rabbit/mouse IgG (Jackson Immu-noResearch) at a 1:3000 dilution and washed in TBST again. Bands were visualized using chemiluminescent substrate (Clarity MaxTM Western ECL Substrate, BIO-RAD).

### Immunofluorescence

Cells were seeded on 15 mm coverslips in a 12-well plate. For fixation, cells were incubated with 4% PFA for 10 min, rinsed 3 times with PBS, permeabilized with 0.2% Triton X-100 for 5 min and rinsed 3 times with PBS. Before staining cells were blocked with 5% BSA in PBS, overnight at 4°C. The next day, cells were stained with primary antibodies: rabbit mAb anti-HLA-A (cat. no. ab52922; Abcam), rabbit pAb anti-HLA-A (cat. no. PA5-29911, Thermo Fisher Scientific) or mouse mAb anti-Flag (cat. no. MA1-91878; Thermo Fisher Scientific) in blocking buffer at 1:500 or 1:1000 dilution at room temperature for 2 h. They were washed 3 times with 1% BSA in PBS and stained with secondary antibodies: Alexa Fluor 488 goat anti-rabbit and Alexa Fluor 555 goat anti-mouse, respectively (Thermo Fisher Scientific) in blocking buffer at 1:2000 dilution, in the dark at room temperature for 1 h, and washed 3 times with 1% BSA in PBS. Nuclei were stained with 300 nM of DAPI (Thermo Fisher Scientific) and samples were mounted using ProLong Diamond Antifade Mountant (Thermo Fisher Scientific). Specimens were imaged using a confocal laser scanning microscope (Leica SP8X, Germany) with a 63x oil immersion lens.

### Proximity ligation assay (PLA)

Cells were pre-treated with respect to fixation, retrieval and permeabilization, and incubated at 37°C in blocking solution for 1 h. Primary antibodies: rabbit mAb anti-HLA-A (cat. no. ab52922; Abcam) and mouse mAb anti-Histone H2B (cat. no. MA5-31410; Invitrogen) were diluted in antibody diluent, applied to samples and incubated at 37°C for 2 h. Cells were washed in a suitable buffer two times for 5 min. The two PLA probes were diluted 1:40 in probe diluent, applied to samples and incubated at 37°C for 60 min. Cells were washed in the appropriate buffer two times for 5 min. Ligation and amplification were performed according to the NaveniFlex protocol (Navinci Diagnostics). Samples were stained with appropriate detection fluorophore (Atto 647N) at 37°Cfor 90 min. Nuclei were stained with 300 nM of DAPI (Thermo Fisher Scientific) and samples were mounted using ProLong Diamond Antifade Mountant (Thermo Fisher Scientific). Specimens were imaged using a confocal laser scanning microscope (Leica SP8X, Germany) with a 63x oil immersion lens.

## References

[1] Neefjes J, Jongsma MLM, Paul P, Bakke O. Towards a systems understanding of MHC class i and MHC class II antigen presentation. Nat Rev Immunol 2011;11:823–36. https://doi.org/10.1038/nri3084.

[2] Kotsias F, Cebrian I, Alloatti A. Antigen processing and presentation. Int Rev Cell Mol Biol 2019;348:69–121. https://doi.org/10.1016/BS.IRCMB.2019.07.005.

[3] von Boehmer H, Kisielow P. Self-nonself discrimination by T cells. Science (1979) 1990;248:1369–73. https://doi.org/10.1126/SCIENCE.1972594.

[4] Ingulli E. Mechanism of cellular rejection in transplantation. Pediatr Nephrol 2010;25:61–74. https://doi.org/10.1007/S00467-008-1020-X.

[5] Sommer S. The importance of immune gene variability (MHC) in evolutionary ecology and conservation. Front Zool 2005;2. https://doi.org/10.1186/1742-9994-2-16.

[6] Wieczorek M, Abualrous ET, Sticht J, Álvaro-Benito M, Stolzenberg S, Noé F, et al. Major Histocompatibility Complex (MHC) Class I and MHC Class II Proteins: Conformational Plasticity in Antigen Presentation. Front Immunol 2017;8. https://doi.org/10.3389/FIMMU.2017.00292.

[7] Raghavan M, del Cid N, Rizvi SM, Peters LR. MHC class I assembly: out and about. Trends Immunol 2008;29:436–43. https://doi.org/10.1016/J.IT.2008.06.004.

[8] Cresswell P, Ackerman AL, Giodini A, Peaper DR, Wearsch PA. Mechanisms of MHC class I-restricted antigen processing and cross-presentation. Immunol Rev 2005;207:145–57. https://doi.org/10.1111/J.0105-2896.2005.00316.X.

[9] Lázaro S, Gamarra D, del Val M. Proteolytic enzymes involved in MHC class I antigen processing: A guerrilla army that partners with the proteasome. Mol Immunol 2015;68:72–6. https://doi.org/10.1016/j.molimm.2015.04.014.

[10] Suh WK, Cohen-Doyle MF, Fruh K, Wang K, Peterson PA, Williams DB. Interaction of MHC class I molecules with the transporter associated with antigen processing. Science (1979) 1994;264:1322–6. https://doi.org/10.1126/science.8191286.

[11] Spiliotis ET, Osorio M, Zúñiga MC, Edidin M. Selective export of MHC class I molecules from the ER after their dissociation from TAP. Immunity 2000;13:841–51. https://doi.org/10.1016/S1074-7613(00)00081-9.

[12] Adiko AC, Babdor J, Gutiérrez-Martínez E, Guermonprez P, Saveanu L. Intracellular transport routes for MHC I and their relevance for antigen cross-presentation. Front Immunol 2015;6:1–11. https://doi.org/10.3389/fimmu.2015.00335.

[13] Yewdell JW, Antón LC, Bennink JR. Defective ribosomal products (DRiPs): a major source of antigenic peptides for MHC class I molecules? The Journal of Immunology 1996;157.

[14] Apcher S, Daskalogianni C, Lejeune F, Manoury B, Imhoos G, Heslop L, et al. Major source of antigenic peptides for the MHC class I pathway is produced during the pioneer round of mRNA translation. Proc Natl Acad Sci U S A 2011;108:11572–7. https://doi.org/10.1073/pnas.1104104108.

[15] Goodenough E, Robinson TM, Zook MB, Flanigan KM, Atkins JF, Howard MT, et al. Cryptic MHC class I-binding peptides are revealed by aminoglycoside-induced stop codon read-through into the 3’ UTR. Proc Natl Acad Sci U S A 2014;111:5670–5. https://doi.org/10.1073/pnas.1402670111.

[16] Laumont CM, Vincent K, Hesnard L, Audemard É, Bonneil É, Laverdure JP, et al. Noncoding regions are the main source of targetable tumor-specific antigens. Sci Transl Med 2018;10. https://doi.org/10.1126/scitranslmed.aau5516.

[17] Weinzierl AO, Maurer D, Altenberend F, Schneiderhan-Marra N, Klingel K, Schoor O, et al. A cryptic vascular endothelial growth factor T-cell epitope: identification and characterization by mass spectrometry and T-cell assays. Cancer Res 2008;68:2447–54. https://doi.org/10.1158/0008-5472.CAN-07-2540.

[18] Malarkannan S, Horng T, Shih PP, Schwab S, Shastri N. Presentation of out-of-frame peptide/MHC class I complexes by a novel translation initiation mechanism. Immunity 1999;10:681–90. https://doi.org/10.1016/S1074-7613(00)80067-9.

[19] Uchihara Y, Permata TBM, Sato H, Kawabata-Iwakawa R, Katada S, Gu W, et al. DNA damage promotes HLA class I presentation by stimulating a pioneer round of translation-associated antigen production. Mol Cell 2022;82:2557-2570.e7. https://doi.org/10.1016/j.molcel.2022.04.030.

[20] Apcher S, Millot G, Daskalogianni C, Scherl A, Manoury B, Fahraeus R. Translation of pre-spliced RNAs in the nuclear compartment generates peptides for the MHC class i pathway. Proc Natl Acad Sci U S A 2013;110:17951–6. https://doi.org/10.1073/pnas.1309956110.

[21] Yewdell JW. MHC Class I Immunopeptidome: Past, Present, and Future. Mol Cell Proteomics 2022;21:100230. https://doi.org/10.1016/J.MCPRO.2022.100230.

## Reference

[1] Suzuki K, Bose P, Leong-Quong RY, Fujita DJ, Riabowol K. REAP: A two minute cell fractionation method. BMC Res Notes 2010;3. https://doi.org/10.1186/1756-0500-3-294.

